# LINE-1 transposable element is expressed in Kaposi’s sarcoma and regulates gene expression in KSHV-infected cells

**DOI:** 10.64898/2026.05.05.722887

**Authors:** Supriya Bhattacharya, Nilabja Roy Chowdhury, Anuj Ahuja, Ayala Fridvald, Boris Rapoport, Vyacheslav Gurevich, Julian Naipauer, Ethel Cesarman, Meir Shamay

**Author notes:** Corresponding author Meir Shamay, **Email:**. These authors contributed equally to this work (Supriya Bhattacharya, Nilabja Roy Chowdhury, Anuj Ahuja).

## Abstract

Long interspersed nuclear element-1 (LINE-1/L1) retrotransposons are expressed in various human cancers. Kaposi’s sarcoma-associated herpesvirus (KSHV) is the etiologic agent of Kaposi’s Sarcoma (KS) and Primary Effusion Lymphoma (PEL), which are characterized by global transcriptional reprogramming. Previously, we have found upregulation of L1 in PEL. Here, we show that knockdown of L1 in PEL cells significantly impaired their proliferation and induced broad transcriptional changes encompassing DNA replication, mitotic progression, and genome maintenance pathways. Analysis of KSHV-infected mouse and human mesenchymal stem cell models revealed that the KS-mimicking microenvironment robustly upregulates L1 expression, with sustained expression observed in KSHV-induced tumor cells even after the loss of the viral episome. Reanalysis of RNA-seq data from KS patient biopsies further showed elevated L1 expression in both endemic and AIDS-associated epidemic KS, with partial reduction following antiretroviral therapy. L1 ORF1p protein expression was confirmed by immunohistochemistry in KS tissues. Reanalysis of the enzymes that can modulate DNA methylation, a process that regulates L1 expression, revealed upregulation of DNMT3A and TET2 in both mouse KSHV tumor models and human KS. Altogether, our findings establish the L1 activation as a hallmark of KSHV-driven oncogenesis.

## Introduction

Kaposi’s sarcoma-associated herpes virus (KSHV/HHV8) is a γ-herpesvirus which is associated with several AIDS-related human malignancies like Kaposi’s sarcoma (KS) ^1^, primary effusion lymphoma (PEL) ^2^, Multicentric Castleman’s disease (MCD) ^3^, as well as the inflammatory disorder KSHV inflammatory cytokine syndrome (KICS) ^4^. KS is the most prevalent form of KSHV-associated disease. There are mainly four forms of KS: Classic, Iatrogenic (following immune suppression), Endemic, and Epidemic. While the endemic form is mainly found in the sub-Saharan African region with HIV-seronegative population, the epidemic form is associated with AIDS ^5^. KSHV-infected spindle cells from KS lesions are poorly differentiated and present gene expression close to endothelial and mesenchymal cells, but different ^6–8^. Epithelial and endothelial cells can go through a transition into mesenchymal cells, a process known as endothelial-to-mesenchymal transition (EndMT). Many cancer cells undergo this transition, leaving them in a transition state between endothelial/epithelial and mesenchymal ^9,10^. KSHV has been shown to induce endothelial cells via EndMT ^11,12^, and mesenchymal cells towards the reverse process (MEndT) ^13,14^. KS spindle cells seem to be in this transition state, leaving the source of these cells, endothelial or mesenchymal, under debate ^15^. PEL is an aggressive form of plasmablastic lymphoma that typically develops in the body cavities of immunocompromised individuals. Cancer development involves the downregulation of the cell’s specific gene expression and the activation of genes resembling those in other tissues, leading to the acquisition of a malignant phenotype ^16,17^. Various epigenetic abnormalities contribute to changes in gene expression, including changes in DNA methylation ^18^, alterations in enhancer switching, microRNA (miRNA) network ^19^, and involvement of transposable elements ^20,21^. PEL cells derived from KSHV-infected patients exhibit a distinctive transcriptional program^22,23^.

Transposable elements (TEs) constitute about 45% of the human genome and continue to shape it. TEs can be classified as DNA transposons, which jump using the encoded transposase, and retrotransposons, which move by a copy-and-paste mechanism via an RNA intermediate. Retrotransposons can be categorized into LTRs (Human endogenous retroviruses or HERVs) and Non-LTRs (LINEs, SINEs, ALUs, VNTRs). Long interspersed nuclear element-1 (LINE-1/L1) is the most prevalent retrotransposon in the human genome, with 500,000 copies that occupies 17% of the human genome ^24^. While most L1 copies have lost their retrotransposition capability, there are still 80-100 copies of evolutionarily young and active L1Hs and L1PA2 that retain the ability to retrotranspose ^25^. L1 transcript encodes two proteins, ORF1p, an RNA-binding protein, and ORF2p, which possess endonuclease and reverse transcriptase activities ^26,27^ and are essential for retrotransposition ^28^.

L1 is tightly regulated by DNA methylation. While DNMT1, DNMT3A, and DNMT3B can add a methyl group to cytosines (DNA methylation), TET1, TET2, and TET3 can reverse this process, leading to DNA demethylation. In cancer cells, both hyper-methylation at specific regulatory regions and global hypo-methylation are often observed. Multiple prior studies have found an elevated L1 ORF1p and ORF2p expression across various cancers^29^ like gastric ^30^, breast ^31^, and pediatric germ layer cancers ^32^. Increased L1 expression might lead to increased retrotransposition and aberrant L1 insertions in new genomic regions, thereby activating oncogenes (e.g., induction of the *Myc* oncogene in breast adenocarcinoma ^21^) and inhibiting tumor suppressor genes (e.g., inhibition of the APC tumor suppressor in colon cancer ^22^). Hypo-methylation and expression of L1 have been detected in Hepatitis C virus (HCV) infection-induced hepatocellular carcinoma (HCC) ^33^, and Hepatitis B virus (HBV) associated HCC ^34^, along with novel L1 retrotranspositions ^35^, and L1 hypo-methylation in serum ^36^. While high L1 expression was also found in Merkel cell polyomavirus (MCPyV)-positive carcinomas^37^, human cytomegalovirus (HCMV) induces L1 expression, and the viral UL44 interacts with L1 ORF2p to promote viral replication ^38^. KSHV-associated PEL cells were shown to enhance L1 retrotransposition^39^. Previously, we reported high L1 expression and reverse-transcriptase activity in KSHV-infected PEL cells ^40^. The importance of this L1 upregulation for PEL cell proliferation was demonstrated by the inhibition of cell growth with RT inhibitors ^40^.

In this study, we investigated the contribution of L1 to PEL proliferation and gene expression by shRNA-mediated L1 knockdown (L1 KD). We found that L1 KD inhibited PEL cell growth. Further RNA-seq analysis of the L1 KD PEL cells revealed enriched pathways involving DNA replication and cell division machineries, supporting the finding of arrested PEL growth. Given that L1 is important for one of the KSHV-associated malignancies, further exploring the transcriptomic profiles of KSHV-infected and non-infected mouse and human mesenchymal stem cells also revealed higher L1 expression in those that were grown in KS-specific medium. RNA-seq analysis from KS patient samples and immunohistochemistry confirmed higher L1 expression in KS tissues. Interestingly, reanalysis of the expression levels of the enzymes that regulate DNA methylation in the same cell models revealed consistent upregulation of both DNMT3A and TET2 in KSHV-associated mouse tumors and human KS.

## Results

### Knockdown of L1 inhibits PEL cell proliferation

Previously, we reported that PEL cells present very high L1 expression, and their growth can be attenuated with RT inhibitors ^40^. Since drugs, including RT inhibitors, can have non-specific effects, we decided to use shRNA to knock down (KD) L1. KSHV-associated PEL (BCBL-1) cells were transduced with shControl or shRNA targeting L1HS (shL1), since it is the youngest and the most active L1 in humans. RNA was isolated and subjected to RT-qPCR using primers against 5’UTR, ORF1, and ORF2 of L1HS (**Fig. 1A**). The KD of L1 was also confirmed by Western blotting with antibodies against L1 ORF1p (**Fig. 1B**). L1 KD led to a significant reduction in BCBL-1 proliferation rate (**Fig. 1C**). The inhibition of cell growth with shRNA targeting L1 supports our previous observation ^40^ that RT inhibitors inhibit cell proliferation. Two cell proliferation markers, CCNA2 and CCNE2 ^41,42^, were also downregulated (**Fig. 1D**). Subsequent RNA-seq revealed a non-significant reduction of total young, intermediate, and old L1 elements, but a significant ∼2.5-fold reduction of the expression specifically of the youngest and most active L1 element, L1HS (**Fig. 1E & F**). The L1PA2, the 2^nd^ most young and active L1 element, also was not decreased significantly. This indicates that L1 KD was specific for L1HS and that the cellular proliferation reduction was mediated specifically due to the decrease in L1HS expression levels.

**Figure 1:**
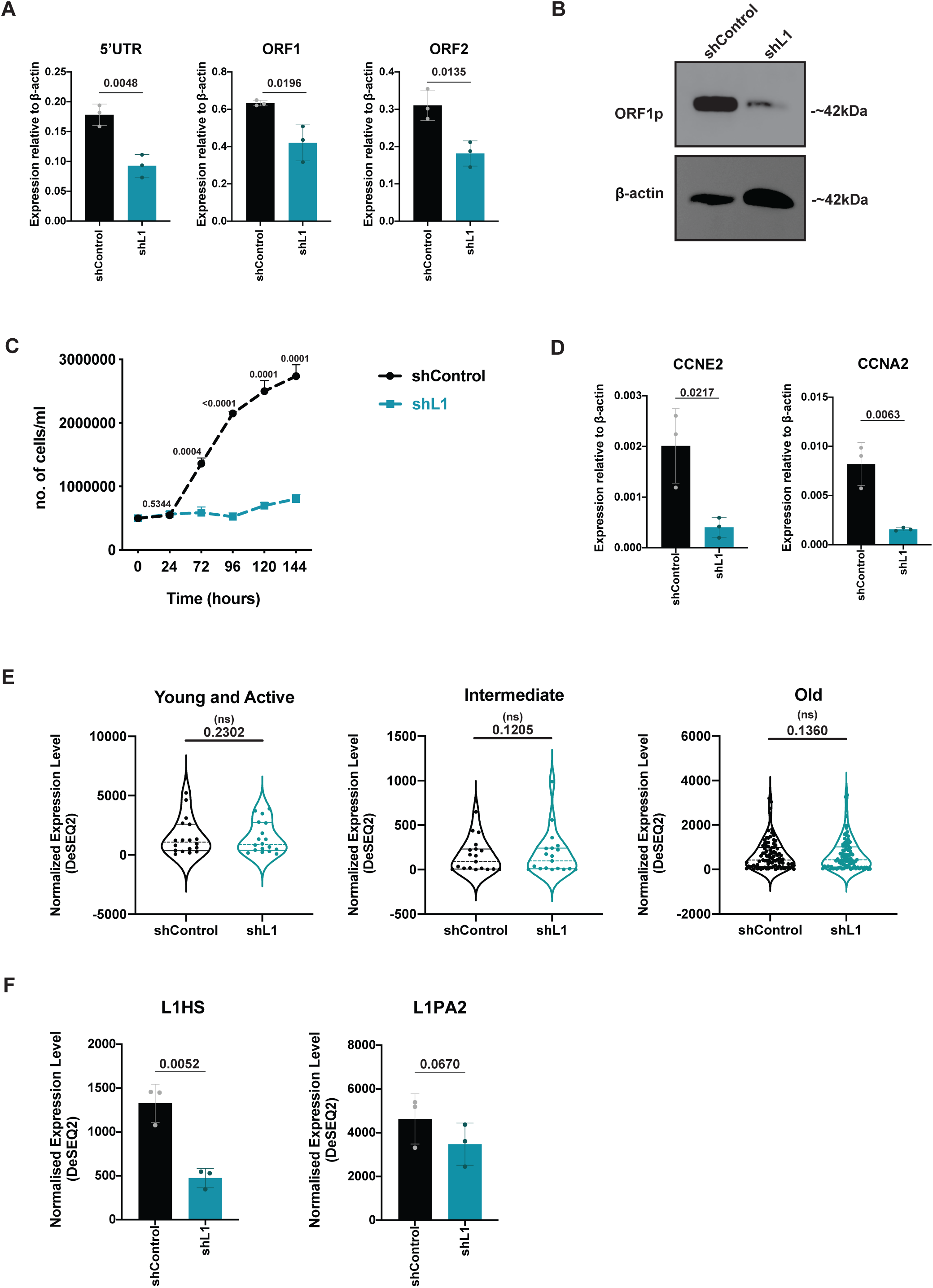
Knockdown of L1 inhibits PEL cell proliferation. (A) RT-qPCR of human L1-specific 5’UTR, ORF1, and ORF2 genes in shControl and shL1 BCBL-1 cells. (B) Western blot of L1 ORF1p and β-Actin control in shControl and shL1 BCBL-1 cells. (C) Cell counts using trypan blue staining ^81^ of control and L1 knockdown cells at respective time-points. (D) RT-qPCR values of CCNE2 and CCNA2 expression levels relative to β-Actin control. (E) Normalized expression levels of Young and Active, Intermediate, and Old L1 elements from RNA-seq data of shControl and shL1 cells are represented with Violin plots. (F) Normalized expression levels of L1HS and L1PA2 elements from the same RNA-seq data of control and knockdown cells. Data represent mean ± S.D. of n = 3. Values in the graph indicate P values from 2-tailed T-tests. (ns) denotes non-significant values.

### L1 contributes to cellular gene expression in KSHV-associated cancer

Next, to gain insights into the effect of L1 KD on the cellular transcriptome, we analyzed the RNA-seq data from shControl and shL1 expressing BCBL-1 cells. This revealed 482 and 977 differentially expressed genes (DEGs) as significantly up- and down-regulated (1.25-fold and FDR-adjusted P value <0.05) in L1 KD cells, respectively (**Fig. 2A**). The top 3 most up- and down-regulated genes were validated using RT-qPCR (**Fig. 2B**). Next, we wanted to get insights into these genes that were significantly differentially expressed. For this purpose, we used Enrichr ^43–45^. Gene Ontology (GO) analysis of the downregulated genes, involving biological processes, showed pathways for DNA metabolic processes, repair, replication, and dsDNA break repair (**Fig. 2C, green bars**). Similar pathways involving DNA replication and cytoskeletal processes, like tubulin and microtubule binding, were seen in the case of GO cellular components (**Fig. 2D, green bars**). For the molecular function involving the downregulated genes, pathways related to the nucleus, mitotic spindle formation, and nuclear chromosome were enriched (**Fig. 2E**). KEGG pathways also showed similar pathways enrichment involving DNA replication, cell cycle, and homologous recombination (**Fig. 2F, green bars**). On the other hand, the upregulated genes were enriched for pathways that involve phagocytic processes (**Fig. 2C-E**). To gain further insights, we selected genes from two pathways, cell cycle (n=128) and DNA replication (n=36) (**Fig. 2G**). While 47 genes that were found to be common among the significantly downregulated genes and the genes that are responsible for cell cycle progression, only 4 genes were common with the significantly upregulated genes. By comparing the DEGs with DNA replication genes, we found 26 genes among the significantly downregulated genes after L1 KD, whereas none were among the upregulated genes. All these observations agree with the growth inhibition of PEL cells after L1 KD (**Fig. 1C**). Altogether, the transcriptomic changes and their enrichment pathways of the shControl and L1 KD BCBL-1 cells indicate that L1 expression is very important for PEL cell proliferation.

**Figure 2:**
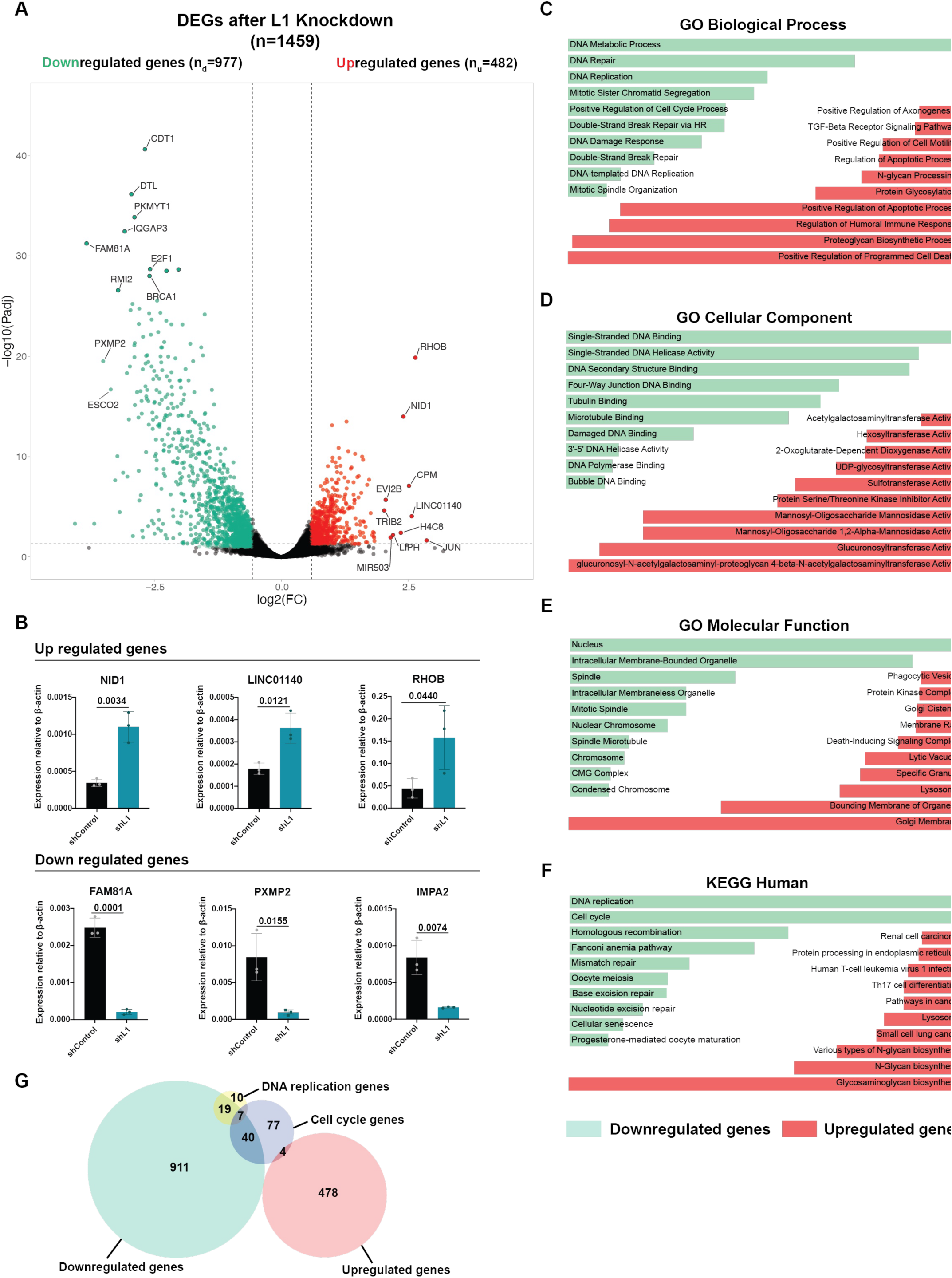
L1 contributes to cellular gene expression in PEL. (A) Volcano plot showing significantly downregulated (marked in green) and upregulated (marked in red) genes after L1 knockdown in BCBL-1 cells. Genes that have an FDR-adjusted P value <0.05 and a fold change of 1.25 were considered significant. The non-significant genes are represented with black colored points. (B) RT-qPCR validation of top 3 up- and down-regulated genes in the L1 knockdown cells. (C-F) Pathway enrichment analyses of significantly down- and up-regulated genes following L1 knockdown of the BCBL-1 cells. Green and red bars indicate the significantly down-regulated and up-regulated genes, respectively. (G) Area-proportional Venn diagram of the significantly up- and down-regulated (green- and salmon-colored circles, respectively) genes with the genes that are responsible for the DNA replication process (yellow circle) and the cell cycle (light blue circle). Data represent mean ± S.D. of n = 3. Values in the graph indicate P values from 2-tailed T-tests.

### L1 is upregulated in mouse MSCs

Given the strong effect of L1 KD on the proliferation of PEL, one of the KSHV-associated malignancies, we hypothesized that in other cells, KSHV infection or its surrounding micro-environment would affect L1 expression as well. Thus, we reanalyzed L1 expression in KSHV-infected mouse mesenchymal stem cells (mMSCs). For that, we chose the RNA-seq dataset from Naipauer et al., 2019 ^46^, where we compared the KSHV- infected mMSCs grown in either a basal medium (**mMSC MEM cells**), which are not tumorigenic, or in a medium mimicking KS-like microenvironment (**mMSC KSM cells**) and the tumors these tumorigenic mMSC KSM cells formed after subcutaneous injection into mice (**mMSC KSM tumors**). This revealed that tumorigenic KSHV-infected mMSC grown in KS-like microenvironment expressed overall more L1 than non-tumorigenic KSHV-infected mMSC grown in basal medium (**Fig. 3B**). This increase was even more profound in the mMSC KSM Tumors. This increase was observed in most L1 elements (**Fig. 3A**) and was more pronounced in the top 10 most-expressed L1 elements (**Fig. 3C**). These results indicate that L1 expression in mMSCs is increased by the proangiogenic medium in which the cells were grown. After that, we reanalyzed our previously published RNA-seq data from the KSHV Hit-and-Run model study ^47^. In this study, KSHV (Bac36)-infected mouse bone-marrow-derived endothelial-lineage cells (tumorigenic **KSHV+ cells**) were either cultured without hygromycin to lose the KSHV episome (non-tumorigenic **KSHV-cells**) or injected into nude mice to form tumors (**KSHV+ tumors**). The tumor cells, when removed from the mice and cultured without hygromycin, lost the episome in vitro, but retained their tumorigenic potential (**Tumorigenic KSHV-cells**). These cells again formed tumors (**KSHV- tumors**) once injected into mice. Reanalysis of the RNA-seq data from these cells revealed that L1 expression once upregulated kept high even following the loss of KSHV (although to a lesser extent), as seen in the case of the tumorigenic KSHV- cells and KSHV- tumors (**Fig. 3D**). The heatmap of all the L1 elements revealed that most of the L1 clusters were upregulated in the KSHV+ tumors (**Fig. 3F**), including the top10 most expressed L1 elements (**Fig. 3E**). The heatmap also revealed that only a small subset of old L1 elements were upregulated in much higher degrees in the Tumorigenic KSHV- cells and KSHV- tumors (like Lx6, L1MC5a, L1MEj, L1M4 etc.), but in the case of KSHV+ tumors, more younger L1s were seen to be upregulated (like L1MdGf, L1MdA, L1MdTf) along with some of the old ones. Altogether, these results indicate that the KS-specific microenvironment preferentially upregulates younger L1s.

**Figure 3:**
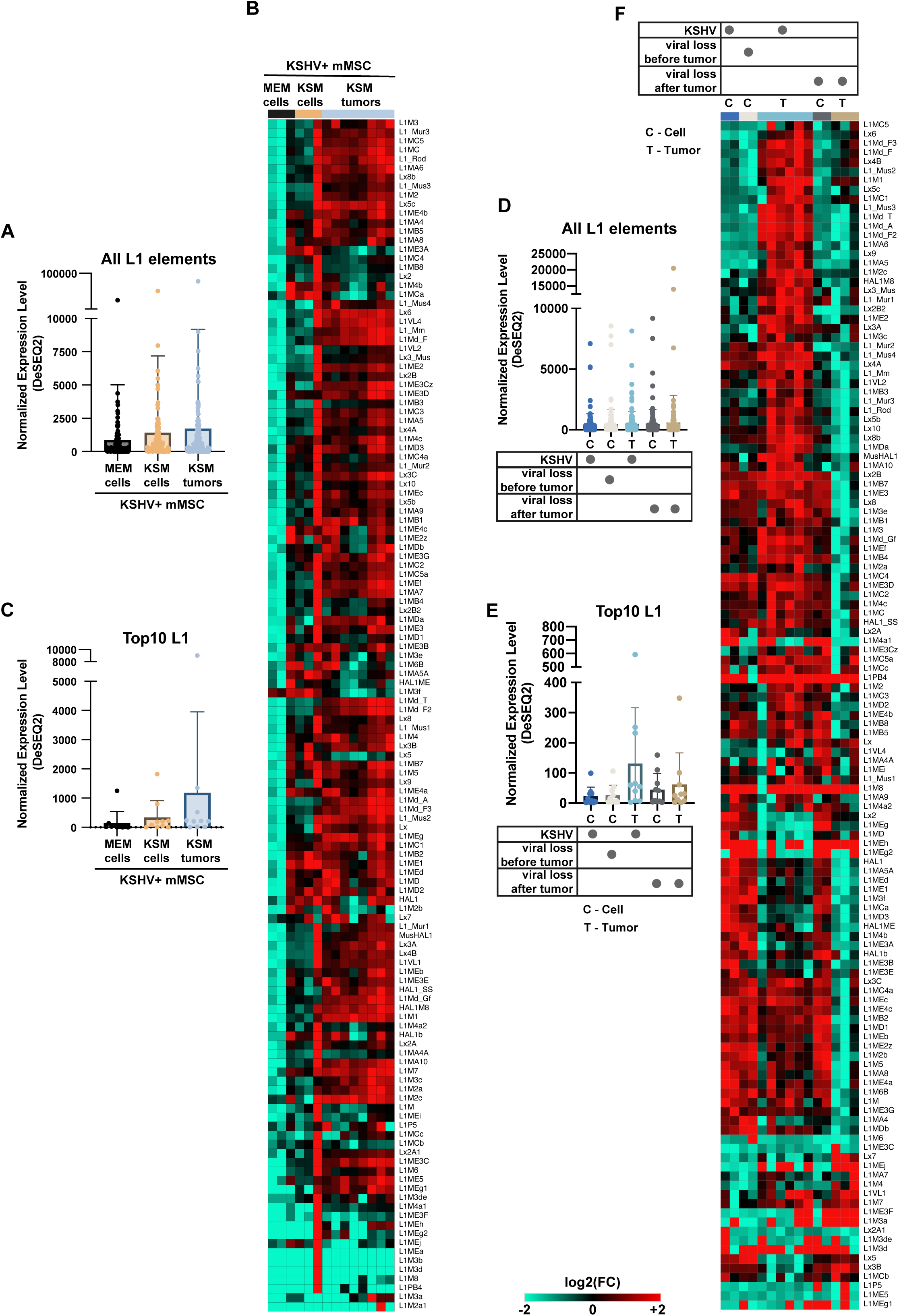
Tumor-specific microenvironment increases L1 expression in mMSCs. (A) Normalized RNA expression levels of all L1 elements from the RNA-seq analysis of KSHV-infected mMSCs grown in MEM, KSM, and the tumors induced by the cells grown in KSM. (B) Heatmap of individual L1 elements from the aforementioned data. (C) Only the top 10 L1 elements from the same data. (D-E) Normalized RNA expression levels of all (D) and top 10 L1 elements (E) from the RNA-seq data of the KSHV ‘Hit & Run’ dataset ^47^. (F) Heatmap of individual L1 elements from the same dataset.

### Upregulation of L1 in human-derived MSCs

Next, we wanted to analyze the L1 expression profile in human bone-marrow-derived mesenchymal stem cells (hMSCs) model for KSHV ^48^. In this study, hMSCs were infected with KSHV (KSHVr.219) and either cultured in basal medium (MEM) or in a medium that mimics a KS-like proangiogenic environment (KSM). The cells were cultured for 72 h, and RNA was subjected to RT-qPCR (using primers for L1HS, the youngest and most active L1 element) and sequencing. RT-qPCR revealed that cells grown in KSM show significantly higher expression of L1 5’UTR, ORF1, and ORF2 than the MEM (**Fig. 4A-C)**. Re-analysis of RNA-seq data also revealed a similar pattern, with most elements being upregulated in cells cultured in the KSM (**Fig. 4D)**. Altogether, this indicates that L1 is overexpressed in hMSCs cells grown in a KS-like microenvironment rather than KSHV infection.

**Figure 4:**
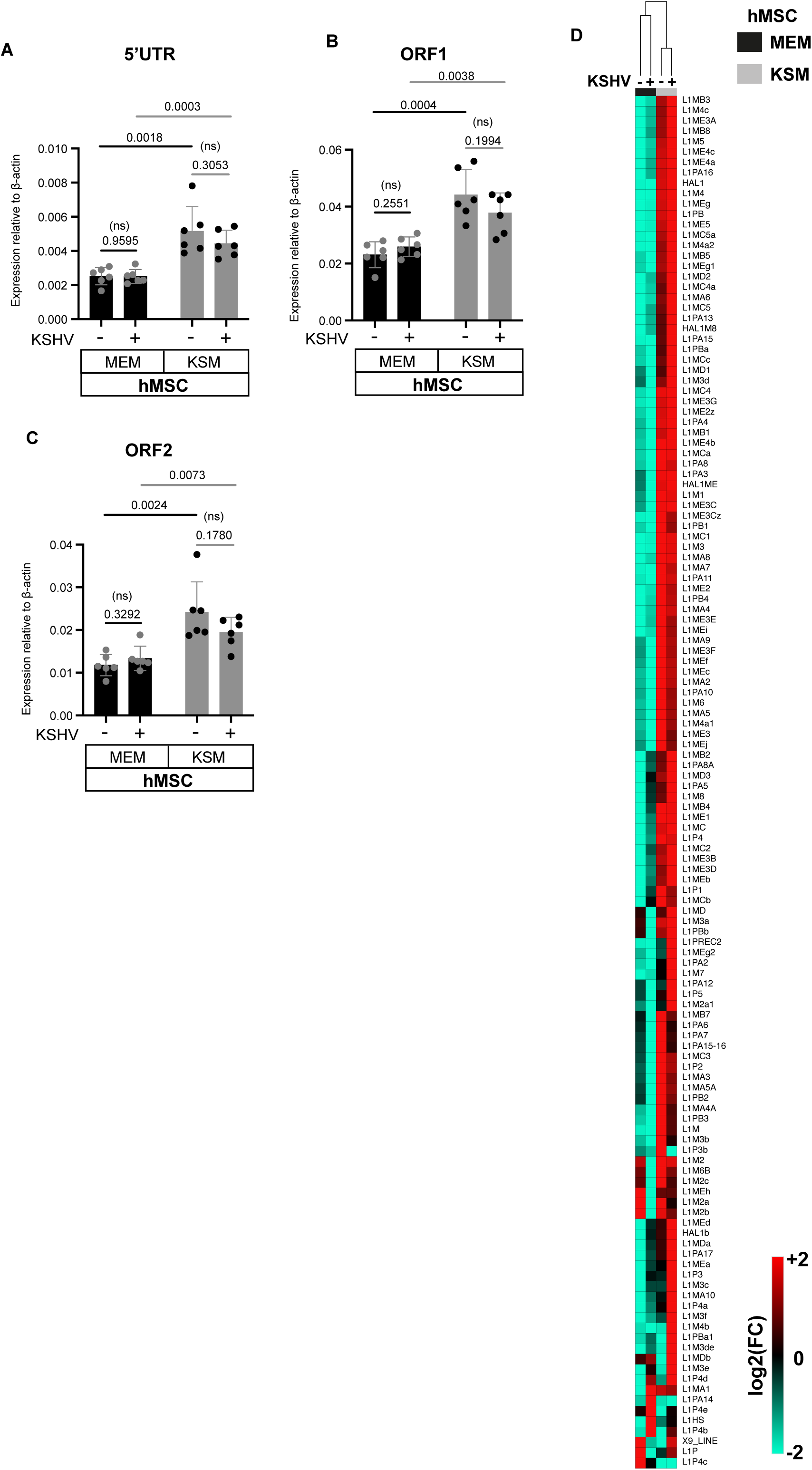
L1 is upregulated in hMSCs due to the KS-specific microenvironment. (A-C) RT-qPCR of human L1-specific 5’UTR (A), ORF1 (B), and ORF2 (C) in hMSCs grown in MEM and KSM. (D) Heatmap of the individual L1 elements from the RNA-seq data ^48^ of the same cells. Data represent mean ± S.D. of n = 6. Values in the graph indicate P values from 2-tailed T-tests. (ns) denotes non-significant values.

### Upregulation of L1 in KS

Since L1 is highly expressed in cells grown in KS-specific medium and in KSHV-induced animal model tumors, we were interested in determining L1 expression in KS. We reanalyzed the RNA-seq data for L1 elements from a large study on KS, including both endemic and epidemic cases ^49^. This revealed upregulation of L1 in KS (**Fig. 5, Supplementary Fig. 1**). Interestingly, the upregulation was much more prominent in the AIDS-associated KS (epidemic) than in the endemic KS. Interestingly, L1 was also highly expressed in the “normal” skin from the same patients with epidemic KS. This L1 upregulation in normal skin was not observed in patients with endemic KS, suggesting that HIV contributes to enhanced L1 expression in epidemic KS. In agreement with this hypothesis, we found reduced L1 expression following antiretroviral treatment (ART) targeting HIV in epidemic KS patients, both in the control skin and KS tissues. This indicates that there is a global change in L1 expression level throughout the epidemic KS patients’ bodies, and ARTs, along with countering HIV, reduce L1 expression in the epidemic KS patients.

**Figure 5:**
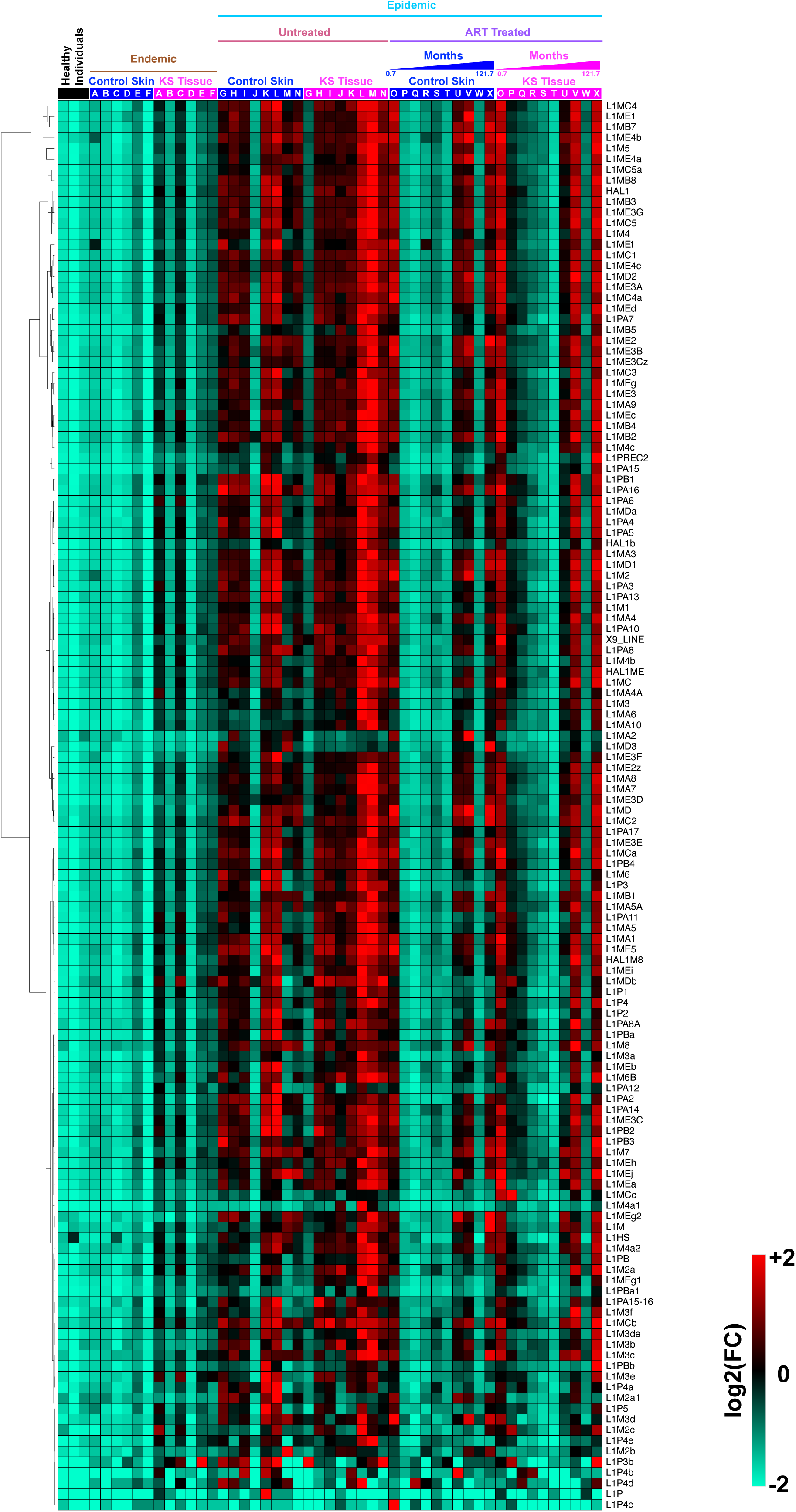
L1 is upregulated in KS tissues. Heatmap representation of individual L1 element expression profile of healthy individuals, endemic and epidemic KS patients, along with their ART-treated counterparts, from Lidenge et al., 2020 ^49^. Individual alphabet in the heatmap denotes individual patients.

To validate this L1 expression in KS, we performed immunohistochemistry (IHC) on skin biopsies from healthy donors and KS for the L1 encoded ORF1 protein (**Fig. 6** and **Supplementary Fig. 2**). This revealed significantly higher expression of L1 in the KS biopsy (**Fig. 6C, D**) compared to healthy skin (**Fig. 6A, B**) from KSHV-negative patients. In summary, we conclude that L1 is highly expressed in KS tissue compared to normal skin.

**Figure 6:**
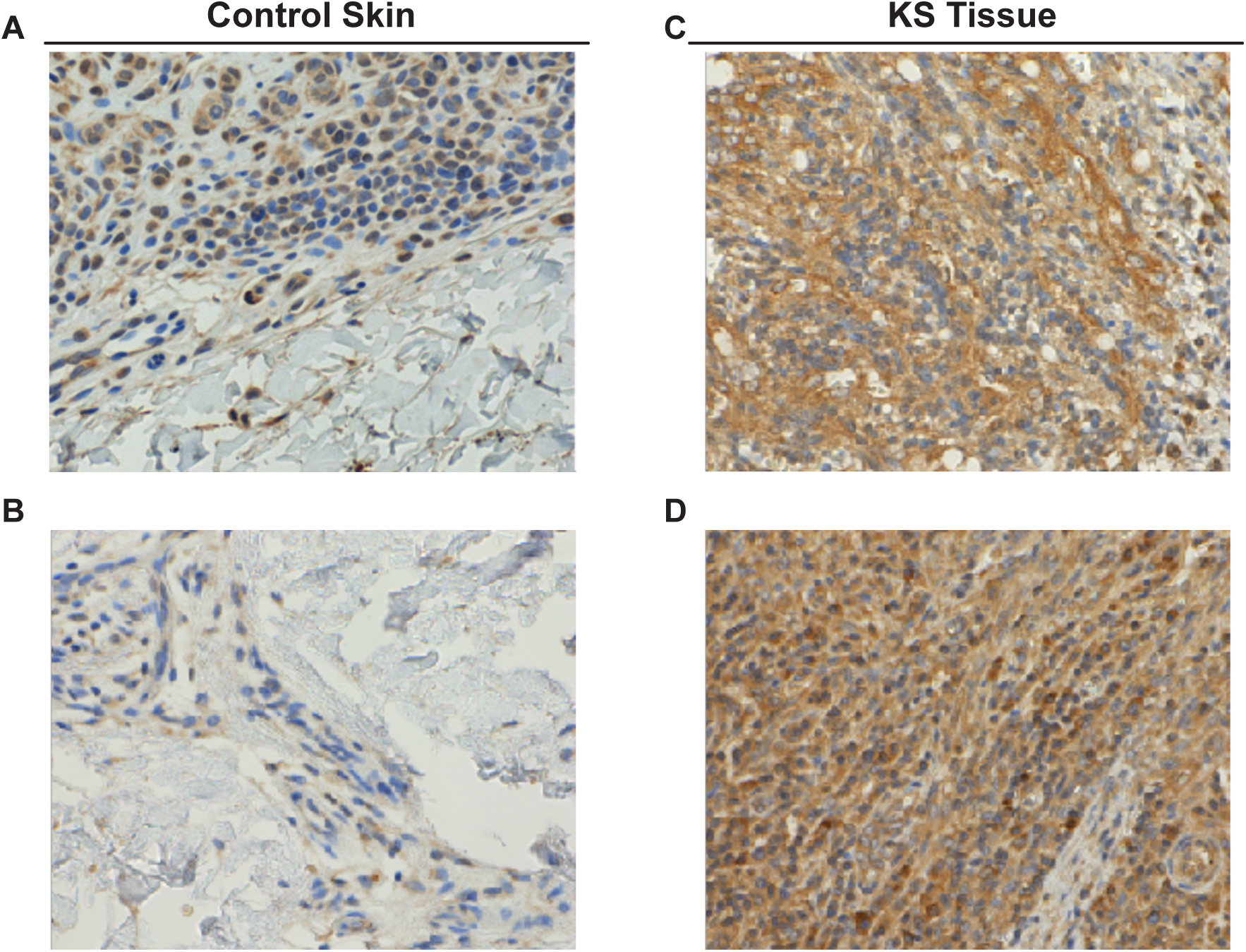
Immunohistochemistry reveals high levels of L1 staining in KS samples. Immunohistochemistry of control skin (A, B) and KS (C, D) tissue samples using L1 ORF1p antibody. Nuclei were stained with hematoxylin stain. (E).

### Upregulation of DNMT3A and TET2 in KS and mouse KS models

The upregulation of L1 in both human KS and the mouse KS models prompted us to search for common epigenetic changes. We specifically searched for the expression of the methylating DNMT1, DNMT3A, and DNMT3B, and the demethylating TET1-3 enzymes, in the same study on both endemic and epidemic KS ^49^. We found elevated levels of DNMT1, DNMT3A, TET1, and TET2 in epidemic KS biopsies compared to skin from healthy individuals (**Fig. 7**). Modest upregulation (not statistically significant) of DNMT1, DNMT3A, TET1, and TET2 was observed in “normal” skin samples from patients with epidemic KS compared to skin from healthy individuals. Therefore, the upregulation of DNMT1, DNMT3A, TET1, and TET2 in epidemic KS was not statistically significant compared with “normal” skin from the same patients, given that these genes were also upregulated in epidemic skin. Elevated levels of DNMT1, DNMT3A, TET1, and TET2 were also observed in epidemic KS following ART treatment. A modest upregulation (not statistically significant) of DNMT3A and TET2 was observed in KS samples from endemic KS.

**Figure 7:**
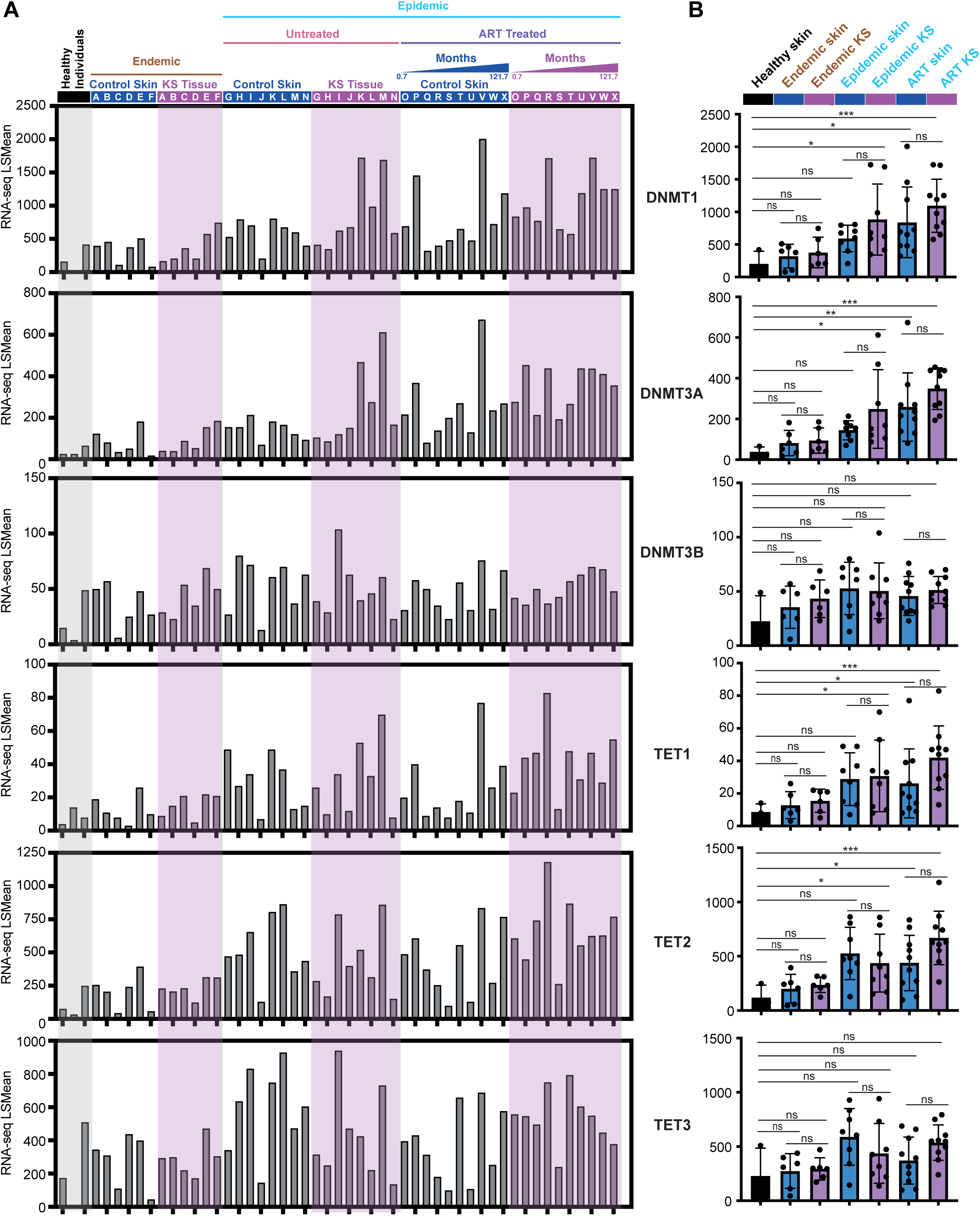
DNMT1, DNMT3A, TET1 and TET2 are upregulated in KS tissues. Graph presenting the expression of DNMT1, DNMT3A, DNMT3B, TET1, TET2, and TET3 in the different samples from Lidenge et al., 2020 ^49^. (A) presents the LSMean reads, (B) presents the LSMean of the different groups inclusinng the SD and P-values from 2-tailed T-tests. (ns) denotes non-significant values.

Reanalysis of RNA-seq from the KSHV-infected mMSCs grown in either a basal medium (**mMSC MEM cells**) or in a medium mimicking KS-like microenvironment (**mMSC KSM cells**) showed no upregulation of DNMT and TET enzymes. However, comparing MEM and KSM cells with tumors formed after subcutaneous injection of KSM cells into mice (**mMSC KSM tumors**), revealed upregulation of Dnmt3a, Tet1, Tet2 and Tet3 (**Fig. 8A, B)**. In addition, we observed downregulation of Dnmt1. Reanalysis of RNA-seq data from the KSHV Hit-and-Run model study ^47^, revealed upregulation of Dnmt3a, Tet2 and Tet3, and downregulation of Dnmt1 in KSHV+ tumors compared to KSHV+ cells (**Fig. 8C, D)**. So, consistent down-regulation of Dnmt1, and upregulation of Dnmt3a, Tet2, and Tet3 was observed in the two KS mouse models. Comparing the tumor mouse models and human KS, revealed shared upregulation of Dnmt3a and Tet2, along with L1 upregulation (**Fig. 8E)**. This led us to the proposed model where both KSHV, microenvironment and HIV infection contribute to upregulation of Dnmt3a and Tet2, increased L1 expression, and development of malignancy (**Fig. 8F)**.

**Figure 8:**
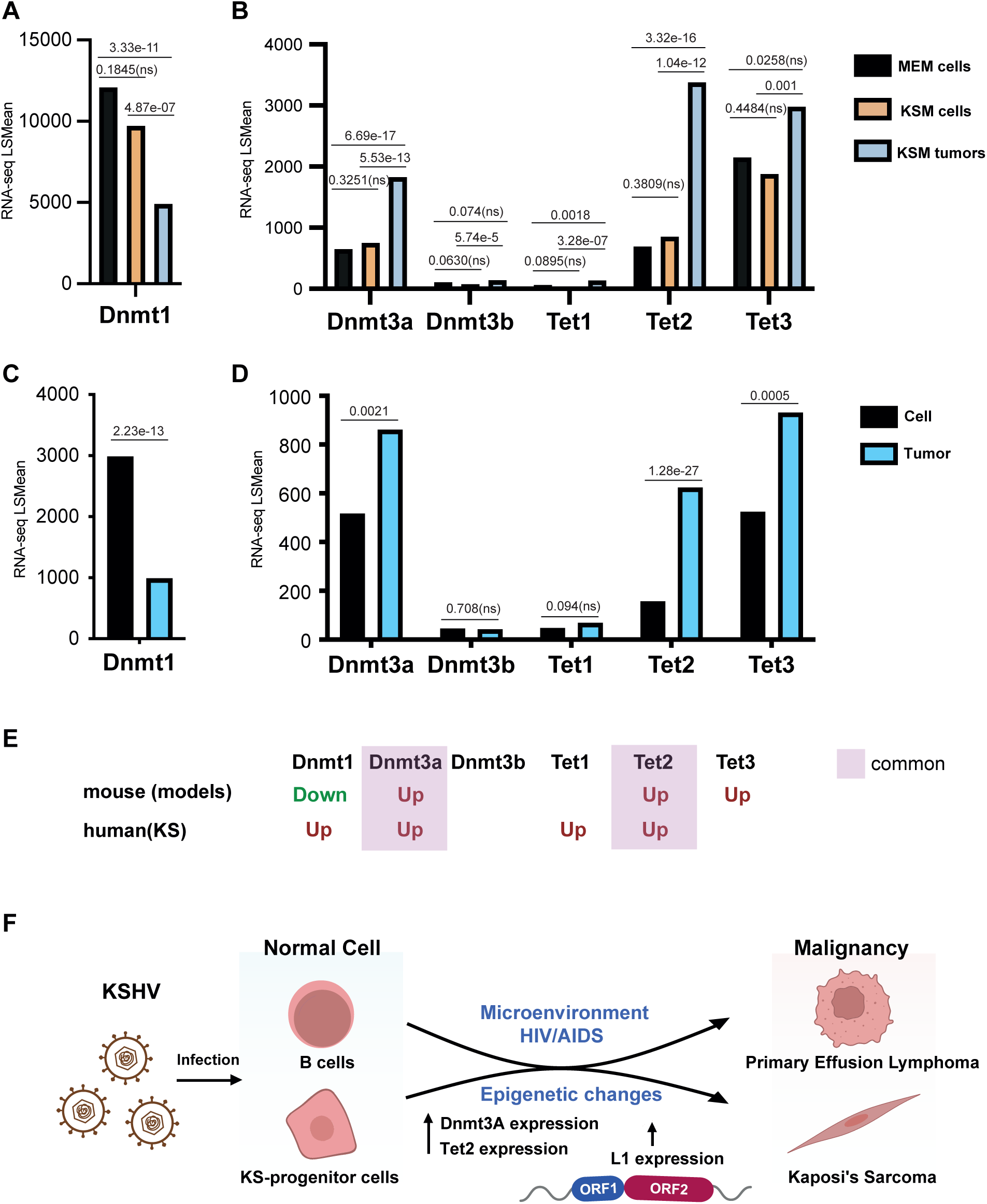
DNMT3A and TET2 are upregulated in both KS mouse tumors and human KS tissues. Graph presenting the expression of DNMT1 (A,C), DNMT3A, DNMT3B, TET1, TET2, and TET3 (B,D) in the different samples from RNA-seq analysis of KSHV-infected mMSCs grown in MEM, KSM, and the tumors induced by the cells grown in KSM (A,B) and the KSHV ‘Hit & Run’ dataset ^47^(C,D). P-values of the RNA-seq via DeSeq2 analysis are presented. (ns) denotes non-significant values. (E) Summary of the RNA-seq gene expression between mouse and human KS, for upregulated (red) and down-regulated (green) enzymes. Common between mouse and human are highlighted (purple). (F) proposed model: KSHV infection in combination with micro-environment and HIV infection leads to upregulation of DNMT3A and TET2, and L1, resulting in the development of PEL and KS.

## Discussion

Oncogenesis leads to global epigenetic changes in infected cells, including altered DNA methylation patterns. L1 is repressed by DNA methylation, and global hypomethylation in cancer leads to upregulated L1 expression across various cancer types. L1 may contribute to tumor growth via transposition-dependent and independent mechanisms. Previously, we reported demethylation of L1 promoters, resulting in higher L1 expression in PEL, one of the KSHV-driven cancers ^40^. We also reported that the usage of RT inhibitors inhibited PEL cell growth. In this study, we provided a multilayered perspective on L1 expression in KSHV-induced oncogenesis, beginning with L1 knockdown in PEL cells and ending with L1 expression in KS tissues, demonstrating that L1 is not merely a bystander in KSHV-induced oncogenesis but an active participant in the proliferative phenotypes of KSHV-associated malignancies.

We hypothesized that L1 might be a contributing factor in PEL growth since treating these cells with RT inhibitors inhibited their growth. Since L1HS is the youngest and most active L1 element, we proceeded with knocking it down using shRNA (L1 KD). This resulted in a significant reduction in PEL proliferation. This aligns with the emerging evidence in other malignancies that L1 has functions beyond insertional mutagenesis ^50,51^. Genes that were downregulated after L1 KD showed interesting pathway enrichments. The molecular processes involve DNA damage repair, replication, mitotic sister chromatid segregation, and cellular component pathways involve ssDNA binding, helicase activity, tubulin and microtubule binding. The molecular function ontology revealed enrichment of pathways involving the nucleus and mitotic spindle formation. All of these are consistent with the observation of arrested PEL cell growth following L1 KD. This also signifies the role of L1 in PEL proliferation.

Tumorigenesis alters the microenvironment, which can impose further epigenetic dysregulation in the tumor itself ^52^. Thus, to assess the effect of the KS microenvironment on L1 expression, we reanalyzed RNA-seq data from mouse and human mesenchymal stem cells (mMSCs and hMSCs, respectively) cultured in either basal medium (MEM) or in KS microenvironment-mimicking medium (KSM) ^46,48^. In both cases, cells grown in KSM presented higher L1 expression. The tumors formed after injecting KSHV-infected mMSCs grown in KSM into mice showed even higher L1 expression. The same phenomenon was observed in the reanalysis of our “Hit and Run” datasets ^53^, in which KSHV(+) tumors show higher expression of most L1 elements. Since full and partial lytic gene expression was observed in these tumors ^53^, we cannot exclude the possibility that lytic induction, either directly or indirectly via a paracrine effect, contributed to higher L1 expression. The finding that L1 expression in KSHV(-) tumor cells following KSHV loss remained high further indicates that L1 expression is part of a tumor-epigenetic memory in these cells. We propose that the ability of KSHV to induce oncogenesis in the absence of HIV in mouse cells can be explained in part by its ability to upregulate L1 in mice.

Reanalysis of RNA-seq data from endemic and epidemic KS samples ^49^ revealed slightly elevated L1 expression in the endemic KS patient samples, while markedly higher L1 expression was detected in the epidemic KS patients. It seems that in HIV-positive KS patients, the control skin also presents higher L1 expression. This suggests that HIV contributes to higher L1 expression in epidemic KS. This idea was further strengthened by the downregulation of L1 following anti-retroviral treatment and the upregulation of L1 expression in patients treated with ART for longer periods, leading to resistance build-up against the therapy ^54–57^. Upregulation of L1 was also verified using IHCs against L1 ORF1p in KS samples, where the spindle-shaped cells in the KS samples signify KSHV-mediated spindle-shaped cell formation. Although sporadic, some L1 staining was found in the healthy control skin; it was limited to the epidermis, which often contains old cells, either deemed to be sloughed off or already dead, which can explain the higher L1 staining ^58^. Besides, the patient’s history only mentioned being KSHV-negative. Other underlying pathologies of inflammatory conditions, like psoriasis ^59^ or systemic lupus erythematosus (SLE), could upregulate L1 expression ^60^, and we could not rule out the control patients’ clinical history of having the aforementioned conditions.

To correlate L1 expression with epigenetic changes in the same cells, we followed the expression of the enzymes that modulate DNA methylation, and found strong upregulation of both the methylating enzyme DNMT3A and the de-methylating TET2. While their activities seem opposing, they are in agreement with DNA methylation in cancer, with specific hyper-methylation at certain promoters and enhancers, and genome-wide hypomethylation. Our previous studies suggested hypermethylation following KSHV infection, and hypomethylation during KSHV-induced tumorigenesis ^53,61,62^. Hypomethylation during tumor development leads to L1 upregulation, so in this respect, the upregulation of TET2 seems most important. In agreement with our finding, higher TET2 and DNMT3A expression were detected during Epstein-Barr virus (EBV) cell transformation ^63,64^. Previously, we have shown that DNMT3A is recruited to specific promoters by LANA ^65^. In the future, it will be interesting to investigate the regulation of TET2 protein recruitment by KSHV.

While L1 was upregulated in both endemic and epidemic KS, its upregulation was higher in epidemic KS, suggesting that HIV contributes to L1 expression. Support for the contribution of HIV to L1 upregulation is its upregulation in “normal” skin from epidemic KS compared to skin from endemic or healthy individuals, and in agreement with a systemic effect via secreted HIV encoded Vpr ^66,67^. In correlation with L1 upregulation, we observed greater upregulation of DNMT3A and TET2 specifically in epidemic KS, suggesting increased epigenetic dysregulation. It is important to note that L1 is upregulated by KSHV and the microenvironment in KS even without HIV (endemic KS), but in combination with AIDS the upregulation is intensified. This L1 increased expression via HIV might explain the increased KS rate in epidemic KS compared to only immunosuppression, as seen in Iatrogenic KS. The inhibition of PEL cell growth following L1 knockdown highlights the dependence of KSHV-associated malignancies on transposable elements and supports L1 inhibition as a potential therapeutic target.

## Methods

### Cell culture

HEK293T cells were maintained in DMEM supplemented with 10% fetal bovine serum (FBS, Gibco), L-glutamine (2 mM, Gibco), Penicillin-streptomycin (100 IU/ml and 100 µg/ml, respectively, Gibco), Sodium-pyruvate (1 mM, Gibco) at 37°C under 5% CO2 atmosphere. BCBL-1 cells were maintained in RPMI-1640 medium with 20% FBS, along with the same supplements. Richard F. Ambinder kindly provided HEK293-T and BCBL-1 cells. The human MSC were cultured in MEM alpha media: supplemented with 20% heat-inactivated fetal bovine serum (FBS) (MSC media); or KS-like proangiogenic growth medium (KS media): Dulbecco’s modified eagle medium supplemented with 30% FBS, 0.2□mg/mL endothelial cell growth factor (ECGF) with 1.2□mg/mL heparin (ReliaTech #300-090H), 0.2□mg/mL endothelial cell growth supplement (ECGS) (Sigma-Aldrich #E0769), insulin/transferrin/selenium (Sigma-Aldrich #I3146), 1% penicillin-streptomycin (Thermo Scientific #15070063), and BME vitamin (MP Biomedicals # ICNA091600449).

### Virus infection

Viral infection was performed with minor modifications to a previously described protocol^48^. Human MSCs were seeded in six-well plates and cultured in MEM alpha medium. After 24 hours, cells were infected with rKSHV.219 at a multiplicity of infection (MOI) of 8. The infection was facilitated by spinoculation in the presence of 8 μg/mL polybrene, by centrifugation at 700 × g for 60 min. Following a 3-hour incubation, the inoculum was removed and replaced with 2 mL of either MEM or KS-like medium in the respective wells.

### L1 Knock-down

HEK293-T cells were transfected with PSPAX2 (Addgene plasmid #12260; https://www.addgene.org/12260/; RRID: Addgene_12260), pMD2.G (Addgene plasmid #12259; https://www.addgene.org/12259/; RRID: Addgene_12259), gifts from Didier Trono, and pVan427 (shControl) and pVan428 (shL1) ^68^, gifts from Gaël Cristofari and Aurélien J. Doucet, (**Supplementary Table 1**), using Polyjet (Signagen, Cat. No. SL100688), to generate lentiviral particles. BCBL-1 cells were transduced with lentiviruses resuspended in RPMI medium supplemented with 8 µg/ml polybrene. The transduced cells were selected with 0.75 µg/ml Puromycin.

### Western Blot

shControl and shL1 BCBL-1 cells were lysed in 0.5 ml RIPA buffer (50 mM Tris [pH 7.4], 150 mM NaCl, 1 mM EDTA, 1% Triton X-100, 0.5% sodium deoxycholate, 0.1% SDS plus protease inhibitor cocktail [Thermo Fisher]). Mouse anti-LINE-1 ORF1p Antibody, clone 4H1 (Merck, Cat. No. MABC 1152), mouse Anti-β-Actin antibody (Merck, A2228), and donkey anti-mouse HRP antibody (Abcam, ab6820) were used in a dilution factor of 1:2500, 1:10000, and 1:5000, respectively, for Western blot analyses.

### Immunohistochemistry

Archived KS biopsies were obtained under Institutional Review Board–approved protocols at Weill Cornell Medicine (New York, NY). No identifying information was associated with samples, and no medical record review was conducted for this study. Immunophenotyping was performed on formalin-fixed, paraffin-embedded tissue sections on a Leica Bond III system using the standard protocol. Sections were pre-treated using heat-mediated antigen retrieval with Sodium-Citrate buffer (pH 6, epitope retrieval solution 1) for 30 mins. The sections were then incubated with monoclonal mouse Anti-LINE-1 ORF1p Antibody, clone 4H1 (EMD Millipore) for 60 min at room temperature and detected using an HRP-conjugated compact polymer system. 3,3’-Diaminobenzidine (DAB) was used as the chromogen. Sections were then counterstained with hematoxylin and mounted with micromount.

### RNA isolation and RT-qPCR

Total RNA from the shControl and shL1 BCBL-1 cells and hMSCs was purified with RNeasy Mini Kit (Qiagen, #74106), according to the manufacturer’s protocol. cDNA synthesis from the RNA was performed using either the High-Capacity cDNA Reverse Transcription Kit (Applied Biosystems, Cat. No. 4374966) or the RevertAid First Strand cDNA Synthesis Kit (Thermo Scientific, #K1622). Subsequently, quantitative PCR (qPCR) was carried out, and the relative expression levels of the genes of interest were determined using target specific primers (**Supplementary Table 2).** Gene expression was calculated using the ΔΔCt ^69^ with β-Actin as the internal control.

### RNA sequencing

Integrity of the isolated RNA from the shControl and shL1 BCBL-1 cells was tested using the Agilent TS HSRNA Kit and Tapestation 4200. 1000ng of total RNA was used for mRNA enrichment with NEBNext mRNA polyA Isolation Module (NEB, #E7490L), and libraries for Illumina sequencing were synthesized with NEBNext Ultra II RNA kit (NEB, #E7770L). The library was quantified using the dsDNA HS Assay kit and Qubit 2.0 (Molecular Probes, Life Technologies). Agilent D1000TS Kit and Tapestation 4200 were used for the subsequent qualification of the quantified library. 500 nM of each library was pooled together and subsequently diluted to 4 nM according to the manufacturer’s instructions. 1.5 pM of library was loaded onto the Flow Cell with 1% PhiX library control. The libraries were sequenced with the Illumina NextSeq 550 platform with single-end reads of 75 cycles.

### Bioinformatic analysis

The FASTq files obtained from the Illumina platforms were first subjected to read quality assessment with FastQC (https://www.bioinformatics.babraham.ac.uk/projects/fastqc/). Any potential residual adapters were trimmed using Trimmomatic ^70^. High-quality reads were used to align with the human genome (hg38) using HISAT2 with default parameters^71^. HT-seq ^72^ was used to count the reads from the BAM alignment files with the annotation from GENCODE, Human Release 49 ^73^. Subsequently, DeSeq2 was used to perform differential gene expression analysis ^74^. Genes showing changes with an FDR-adjusted P<0.05 were considered for further downstream analyses and experimental conclusions. Volcano plot of the DEGs was made using ggPlot2. An area-proportional venn diagram was made using DeepVenn ^75^.

For LINE-1 (L1) expression analysis, the FASTq files were aligned to the human reference genome (hg38) using the STAR aligner (v2.7.11b) ^76^. To focus on uniquely mapped reads, **-outFilterMultimapNmax** parameter was set to **1** which allowed the exclusion of sequences aligning to multiple loci. Reads with the highest alignment score in case of multiple alignments were retained using the parameter **- outFilterMultimapScoreRange 1**. Alignment quality was ensured by applying **- outFilterMismatchNoverLmax** to **0.03** to apply a mismatch threshold of 3% per base. L1 elements were annotated using Repeatmasker ^77^ and quantified using featureCounts ^78^, which was normalized using DeSeq2 ^74^, scaling the mapped reads to the hg38 reference genome in a replicate-wise manner. Further classification of L1 elements based on evolutionary age (young and active, intermediate, and old) was performed based on previously reported criteria ^79,80^. Bar graphs and violin plots were made using GraphPad Prism software (v10). Heatmaps of individual L1 elements were created using Morpheus (https://software.broadinstitute.org/morpheus/).

### Quantification and statistical analyses

GraphPad Prism (v9.5.1), unless stated otherwise, was used to quantify data, generate graphs, and perform statistical analyses with 2-tailed Student’s t-tests.

### Data availability

The RNAseq data that were reanalyzed for L1 expression are publicly available in repositories with accession nos. as follows: GSE141868 ^46^, GSE144101 ^47^, GSE260925 ^48^, and GSE147704 ^49^. The raw RNA sequencing data of the control and L1 KD cells were uploaded to NCBI SRA and will be made publicly available upon acceptance of this manuscript.

## Acknowledgments

We would like to thank Didier Trono, Gaël Cristofari, and Aurélien J. Doucet for kindly providing the plasmids, Richard F. Ambinder and S. Diane Hayward for cell lines. Immunohistochemistry for this study was performed by the Center for Translational Pathology at Weill Cornell Medicine.

## Author Contributions

Conceived and designed the experiments: S.B., N.R.C., A.A., V.G., E.C., J.N., M.S. Generated tools & reagents: S.B., N.R.C., A.A., V.G. Performed the experiments: S.B., N.R.C., A.A., A.F. Analyzed the RT-qPCR data: S.B., N.R.C., A.A., A.F. Bioinformatic analysis: N.R.C., M.S. Immunohistochemistry: A.A., E.C. Wrote the paper: N.R.C., M.S.

## Supplementary figure legends

**Supplementary Figure 1**: **Individual L1 count from healthy and KS patient biopsies.** RNA-seq analysis from the study by Lidenge et al. 2020 was performed on L1 transposable elements. Individual L1 elements: L1PB, L1M6, L1M3de, L1M4a1, L1M2a, L1PREC2, L1M, HAL1b are presented for healthy controls (n = 3), skin and KS from endemic (n = 6), skin and KS from epidemic (n = 8), and skin and KS following ART treatment (n = 10). Data represent mean ± S.D. Values in the graph indicate P values from 2-tailed T-tests. (ns) denotes non-significant values.

**Supplementary Figure 2: Detection of L1 ORF1p in KS.** (A-B) L1 ORF1p immunohistochemistry of control skin (A) and KS tissue samples (B) reveals significantly more expression of L1. Spindle-shaped cells present in the KS samples and not in the control skin. Cell nuclei were stained with hematoxylin.

