## Supplementary figures for "LINE-1 transposable element is expressed in Kaposi’s sarcoma and regulates gene expression in KSHV-infected cells"

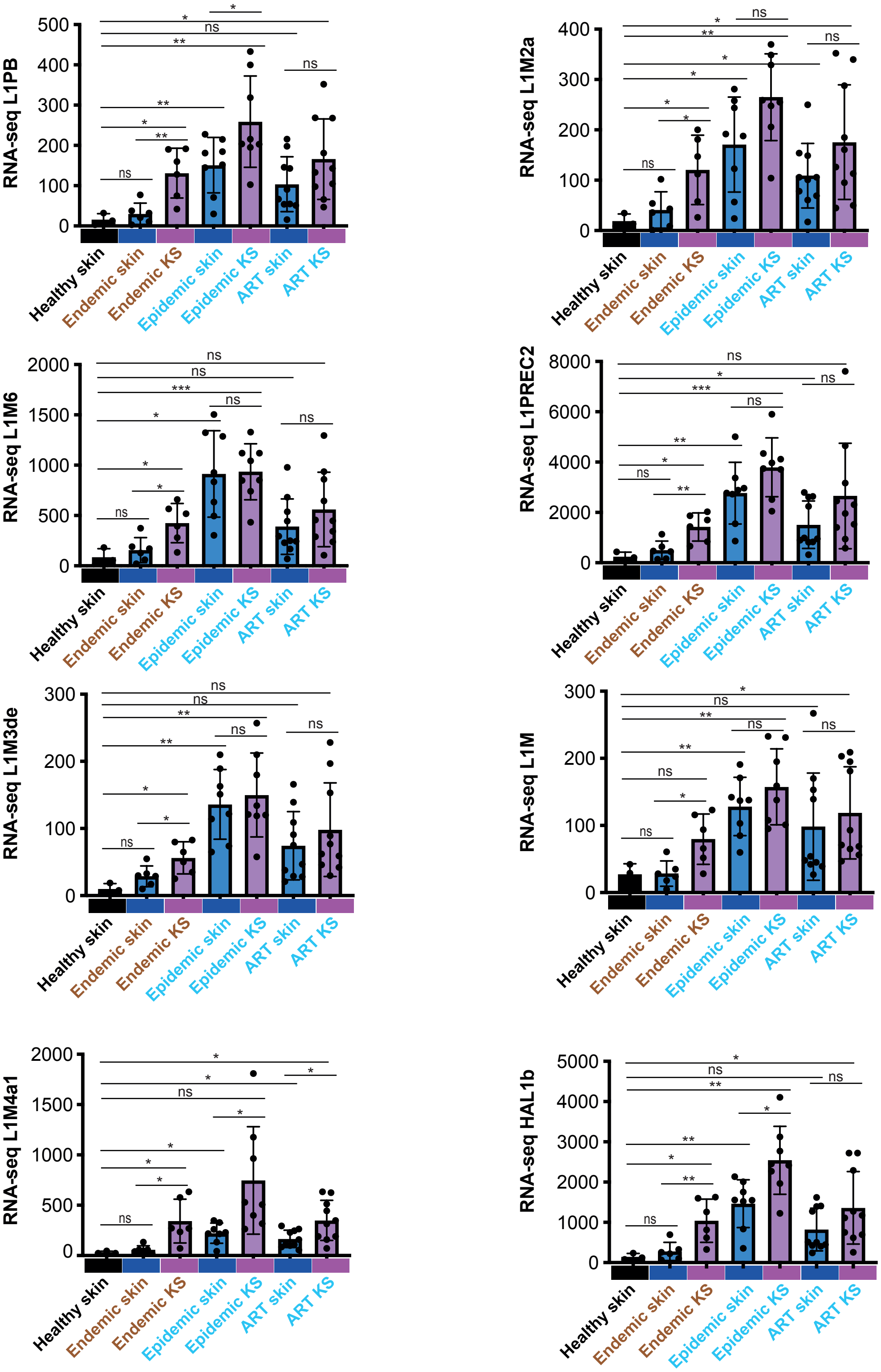


**Supplementary Figure 1**: Individual L1 count from healthy and KS patient biopsies. RNA-seq analysis from the study by Lidenge et al. 2020 was performed on L1 transposable elements. Individual L1 elements: L1PB, L1M6, L1M3de, L1M4a1, L1M2a, L1PREC2, L1M, HAL1b are presented for healthy controls (n = 3), skin and KS from endemic (n = 6), skin and KS from epidemic (n = 8), and skin and KS folowing ART treatment (n = 10). Data represent mean ± S.D. Values in the graph indicate P values from 2-tailed T-tests. (ns) denotes non-significant values.

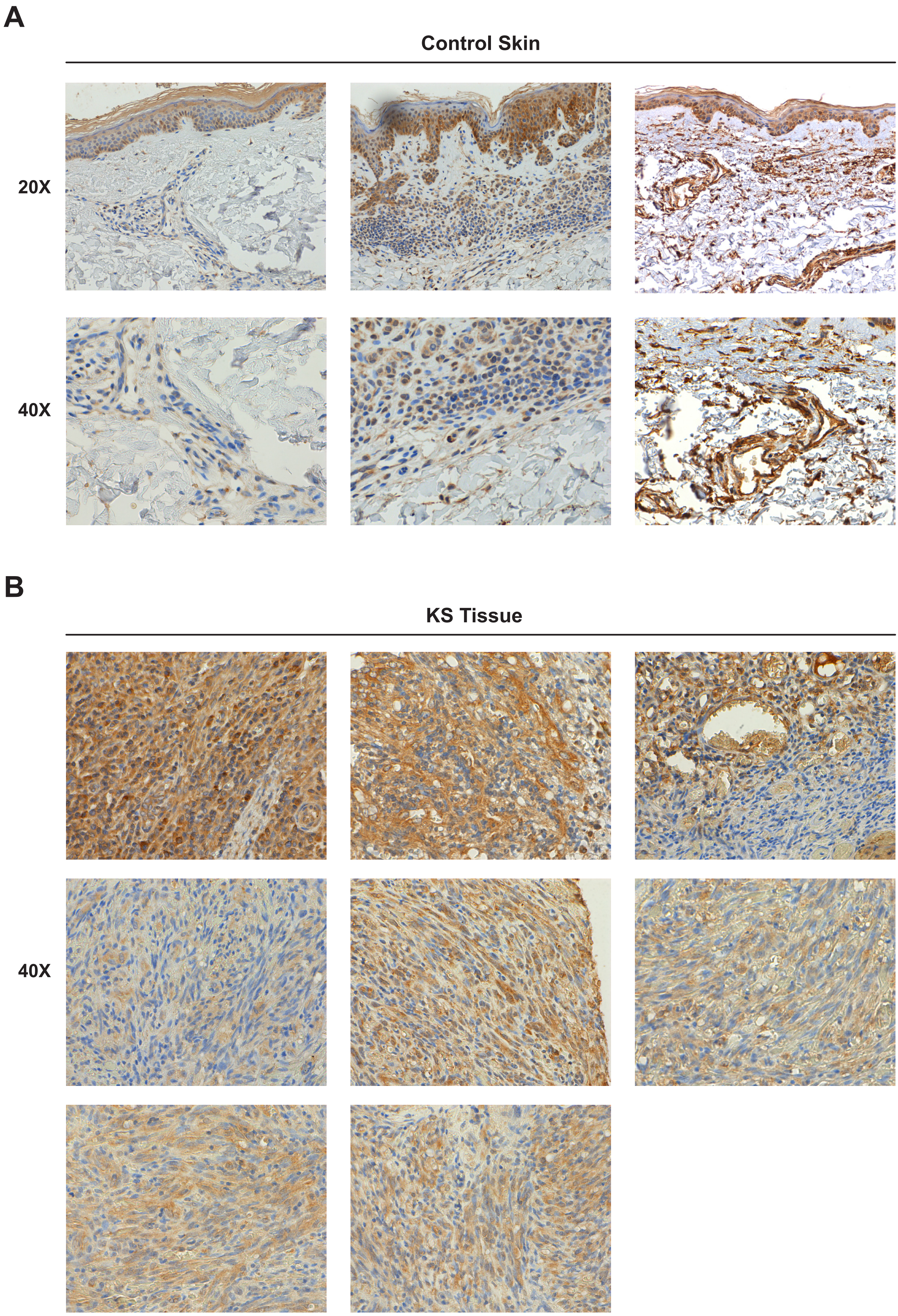
A
Control Skin
20X
40X
B
KS Tissue
40X

**Supplementary Figure 2: Detection of L1 ORF1p in KS.** (A-B) L1 ORF1p immunohistochemistry of control skin (A) and KS tissue samples (B) reveals significantly more expression of L1. Spindle-shaped cells present in the KS samples and not in the control skin. Cell nuclei were stained with hematoxylin.
